# Role of brown adipose tissue-specific sympathetic and sensory innervations in menthol-induced energy-expending phenotype

**DOI:** 10.64898/2026.01.07.698109

**Authors:** Roshan Lal, Neha Soni, Kushhagra Agarwal, Kanthi Kiran Kondepudi, Pragyanshu Khare, Katharina Zimmermann, Małgorzata Karbownik-Lewińska, Adam Gesing, Vaishali Aggarwal, Kanwaljit Chopra, Mahendra Bishnoi

## Abstract

Menthol, a TRPM8 agonist and pharmacological cold mimic, activates brown adipose tissue (BAT) to increase adaptive thermogenesis and whole-body energy expenditure. BAT is highly innervated with sensory and sympathetic nerve fibers. There is existing literature that sympathetic nerves have a critical role in regulating menthol’s (or cold) effect on energy homeostasis, but there is minimal literature about the involvement of somatosensory innervations. Here, we ought to investigate whether short-term topical application (3 days) of menthol elicits sympathetic and sensory nerves. We used chemical denervation models of complete (100 mg/kg; 6-OHDA and 125 mg/kg; capsaicin, s.c.) and localized (in BAT, 20 µL of 6-OHDA (20 mg/ml) and 20µL of capsaicin (20µg/µL) in each BAT lobe) ablation of sympathetic (6-OHDA) and sensory (capsaicin) innervations to explore their roles in menthol induced BAT activation and energy expanding phenotype. In the present study we have shown that (i) short-term topical application of menthol (10 % menthol for 3 consecutive days) elicits sympathetic and sensory innervations, and induces thermogenesis, lipolysis and mitochondrial biogenesis in mice BAT; (ii) localized ablation of sympathetic innervation prevented menthol induced BAT activation and lipolysis (iii) localized ablation of sensory neurons augments sympathetic innervations induced BAT activation and modulated thermoregulation.

Collectively, our results suggest that BAT activation following short-term topical application of menthol is dependent primarily but not exclusively on centrally mediated SNS activation. TRPV1-positive sensory neurons are partially necessary for thermoregulation and efficient BAT activation.

## 1. Introduction

Increasing whole-body energy expenditure has emerged as a promising strategy to combat obesity and related metabolic disorders [1]. Mammals possess three types of adipose tissue: white adipose tissue (WAT), specialized for energy storage; brown adipose tissue (BAT), specialized for energy dissipation via thermogenesis; and the recently characterized “brite” (brown-in-white) adipocytes, which are inducible thermogenic cells within WAT depots [2]. BAT plays a central role in non-shivering thermogenesis and energy homeostasis, making it a key target for obesity intervention strategies [3–5]. While cold exposure is a potent activator of BAT, pharmacological cold mimetics such as menthol and icilin also stimulate BAT thermogenesis and increase energy expenditure in mice [6–8].

Menthol activates Transient Receptor Potential Melastatin-8 (TRPM8), a well-characterized cold sensor highly expressed in peripheral sensory neurons and in both white and brown adipocytes [6, 9–12]. Topical menthol application stimulates TRPM8, triggering behavioral and physiological heat-gain responses such as increased oxygen consumption, non-shivering thermogenesis, and elevated core body temperature (CBT) [13, 14]. These effects are mediated via sympathetic nervous system (SNS) activation, leading to upregulation of uncoupling protein 1 (UCP1) and increased norepinephrine levels in BAT [8]. In diet-induced obese mice, topical menthol significantly increases basal metabolic rate without altering food intake [15, 16]. Moreover, intermittent application over two weeks reduces WAT mass and body weight through sustained hyperthermia [17]. Prior work from our laboratory has shown that both oral and topical menthol induce BAT thermogenesis, promote WAT browning, and activate glucagon signaling in high-fat diet (HFD)-fed mice [18]. Additionally, menthol induces a mitochondrial and energy-expending phenotype in 3T3-L1 adipocytes [19]. Most recently, we demonstrated that topical menthol enhances adaptive thermogenesis, warm-seeking behavior, lipid oxidation, and sympathetic innervation in BAT [20]. Dietary menthol at higher concentration (0.5 %) significantly reduced food intake, body weight, and visceral fat [21]. In addition, at lower concentrations (0.25 %), menthol improved autonomic thermoregulation by activating TRPM8 without significantly affecting food intake or body weight [21].

BAT is richly innervated by both sympathetic and sensory nerve fibers, each contributing to its thermogenic regulation [22–24]. The role of sympathetic innervation is well established: norepinephrine or epinephrine (NE or EPI), co-released with neuropeptide Y (NPY) and adenosine triphosphate (ATP) from sympathetic terminals, regulates thermogenesis and lipid metabolism [25, 26]. Chemical or surgical sympathectomy of interscapular BAT drastically reduces UCP1 expression, mitochondrial content, and glucose uptake—all of which can be restored by exogenous NE [27–29]. In contrast, the role of sensory innervation in BAT function remains less defined. Emerging evidence suggests sensory nerves provide feedback to sympathetic signaling and contribute to thermogenic regulation. Sensory neurons may respond to temperature changes or lipolytic byproducts by releasing neuropeptides such as calcitonin gene-related peptide (CGRP) and substance P (SP), which modulate adipose metabolism [30, 31]. Global sensory denervation with capsaicin impairs BAT mass and UCP1 expression in cold-exposed animals [32, 33], and recent studies implicate sensory afferents as essential for WAT browning, independent of sympathetic input [24]. Notably, PIEZO2-expressing sensory neurons have been shown to inhibit sympathetic outflow and thus regulate adipose function [34–36].

Despite these insights, no prior study has directly examined how sensory afferents contribute to BAT activation by pharmacological cold mimetics. To address this gap, our study aimed to (1) determine whether short-term (3-day) topical menthol application induces both sympathetic and sensory innervation while stimulating BAT thermogenesis, and (2) delineate the individual contributions of sensory and sympathetic nerves using selective chemical denervation—6-hydroxydopamine (6-OHDA) for sympathetic and capsaicin for sensory ablation. Collectively, our findings reveal a previously unrecognized, yet essential, role of BAT sensory innervation in mediating menthol-induced thermogenesis, highlighting its therapeutic potential in cold-mimetic strategies against obesity.

## 2. Methodology

### 2.1 Animals

Male C57BL/6J (22-25 g; 8-10 weeks) acquired from IMTech Center for Animal Resources and Experimentation (iCARE), Chandigarh, India, and housed in Animal Experimentation facility (AEF) at National Agri-Food Biomanufacturing Institute (NABI), Mohali, India. All the mice were maintained under a pathogen-free environment under controlled temperature (25±2°C), humidity conditions (55±5%), and a 12 h light-dark cycle. Mice were provided with diet and water ad libitum during the experiment. All animal experiments were approved by IAEC, NABI (approval no. NABI/2039/CPCSEA/IAEC/2024/16), and experiments were conducted in accordance with the Committee for the Purpose of Control and Supervision of Experiments on Animals (CCSEA) guidelines on the use and care of experimental animals.

### 2.2 Experimental protocol

#### 2.2.1 Short-term menthol application

Briefly, mice were acclimatized to experimental conditions and randomized into two groups (N=8/group), namely (i) vehicle control (NPD, 4g/kg vehicle); (ii) menthol (NPD, 4g/kg 10% menthol). Topical menthol was applied for 3 days post-acclimatization (single application/ day). All applications were performed on tail-restrained mice on the unshaved dorsal surface from the interscapular area to the sacral area.

#### 2.2.2 Chemical denervation of SNS with 6-OHDA

Complete peripheral sympathectomy was achieved by injection (100 mg/kg; i.p.) of the catecholaminergic neurotoxin, 6-OHDA. Mice were left for forty-eight hours after injection [8]. For local iBAT sympathectomy interscapular BAT depots were exposed by an incision. 6-OHDA (10 mg/mL) was dissolved in 0.15 mol/l NaCl and 1% ascorbic acid solution. 6-OHDA solution, at a dose of 100 µg/BAT lobe, was bilaterally injected into BAT lobes using a Hamilton syringe. Mice were allowed to recover for 7 days before short-term menthol application [24, 37].

Mice were randomly divided into six groups (i) vehicle (VEH) (ii) menthol (MTH) (iii) local BAT SNS denervation (L-6OHDA) with no menthol application (iv) local BAT SNS denervation with short-term topical menthol application (MTH +L-6OHDA) (v) complete body SNS-denervation (C-6OHDA) (vi) complete body SNS-denervation with short-term topical menthol application (MTH + C-OHDA). Topical menthol was applied for 3 days (single application/per day) in the respective groups, and CBT measurement, BAT thermography, and cold tolerance test were performed. After completing phenotypic assessments, animals were sacrificed, serum, BAT, and hypothalamus were collected and stored at -80°C for further analysis.

#### 2.2.3 Chemical (capsaicin) induced sensory denervation

Complete peripheral sensory denervation was achieved with capsaicin administered into subcutaneous fat tissues of the neck in two injections (50 mg/kg and 75 mg/kg), 24 h apart under isoflurane anaesthesia. Atropine (5 mg/kg, i.p.) was administered immediately before capsaicin injection to prevent acute cardiopulmonary effects of excessively released sensory mediators. Local sensory denervation of interscapular BAT was achieved as previously described [38]. Briefly, 10µl Hamilton syringe was used to deliver 10 microinjections (2 μl/injection) of 20 μg/μl of capsaicin (1:10, ethanol: olive oil) [38]. After achieving the ablation of sensory nerves globally and in local BAT tissue, mice were randomized into six groups (i) VEH (ii) MTH (iii) local BAT sensory denervation (L-CAP) with no MTH application (iv) local BAT sensory-denervation with short-term topical menthol application (MTH +L-CAP) (v) complete body sensory-denervation (C-CAP) (vi) complete body sensory-denervation with short-term topical menthol application (MTH+ C-CAP). Topical application of menthol was done for 3 days. CBT, BAT heat production, and cold tolerance test were measured after the last application of menthol. Next, mice were sacrificed, and serum, BAT, and hypothalamus were harvested and stored at -80°C for ELISA and gene expression analysis.

### 2.3 Oral glucose tolerance test (OGTT)

OGTT was performed in 12 h fasted mice. Briefly, fasted mice were administered 2 g/kg BW glucose orally. Blood glucose will be measured at 0 (before administration),15, 30, 45, 60, and 120 min, using Glucocard™ (Arkray Factory Inc., Shiga, Japan) [39].

### 2.4 Hot plate test

Mice were placed on hot plate surface with temperature maintained at 55±1 °C. The time latency for paw licking or jumping was noted as first sign to avoid heat was recorded as pain threshold. To avoid any paw tissue damage cut-off time was 25 s [40].

### 2.5 CBT

CBT was measured using a rectal probe (Orchid Scientific and Innovative India Pvt. Ltd). Briefly, temperature probe is inserted 10 mm in mice rectum. Temperature readings were taken at 0 (before menthol application), 30, 60, 120, and 180 min post-menthol application. Temperature probe was wiped with 70 % ethanol between the readings [41].

### 2.6 Infrared thermography

Thermogenic activity of the iBAT was assessed by infrared thermography. Mice housed at ambient room temperature (25 ± 2°C) were treated with topical menthol. Approximately 10 pictures of each mouse were taken at 0, 30, 60,120, 180, and 240 min post-menthol application (FLIR T530 thermal camera, with emissivity set on 0.98, standardized focal length, using rainbow contrast palette). Two standardized regions of interest, ROI-1 (nuchal/neck region) and ROI-2 (interscapular), were chosen, and temperature difference between ROI-1(nuchal/neck region) to ROI-2 (interscapular) was calculated. FLIR tools app was used to analyse the images [20].

### 2.7 Cold tolerance test

16 h overnight fasted mice were individually housed at 4°C for 4h with free access to water. Rectal temperature was measured every hour during cold exposure using a rectal probe [42].

### 2.8 Gene expression analysis

Briefly, total RNA from BAT and hypothalamus was extracted using the Trizol-chloroform-isopropyl alcohol method previously described [43]. After quantification and quality check, DNase treatment was given for removal of any contamination of genomic DNA. RNA was then transcribed into cDNA using revertaid cDNA synthesis kit (Cat. No. K1622, Thermo Fisher Scientific, United States). The synthesised cDNA was used for gene expression analysis. The qRT-PCR was performed on CFX96 touch real-time PCR detection system (Bio-Rad Laboratories, United States) using SYBR green supermix (Cat. No. K1622, Bio-Rad Laboratories, United States) using these conditions: 95°C for 2 minutes (initial denaturation), followed by (denaturation at 95 °C for 5 second, annealing and elongation at 60°C for 30 second) × 40 cycles, final extension at 60°C for 5 minute and melt curve analysis between 60-95°C with 0.5°C/5 second increment. Analysis of relative change in gene expression will be done using the 2^−ΔΔct^ method [44]. Ct values were normalized to TBP (housekeeping) gene, and values were expressed as fold change with reference to vehicle control. The list of primers is given in **Table 1**.

### 2.9 Enzyme-linked immunosorbent assay (ELISA) estimations

Serum was isolated from blood. BAT samples were homogenized in ice-cold phosphate-buffered saline to prepare tissue lysates. EPI (Cat. No.-E-EL-0045, Elabscience Biotechnology Inc.); NPY (Cat. No.-E0704Mo, Shanghai Korain Biotech Co. Ltd.); ATP (Cat. No.-E0665Mo, Shanghai Korain Biotech Co. Ltd., CGRP (Cat. No.-E0556Mo, Shanghai Korain Biotech Co. Ltd.) SP (Cat. No.-E0357Mo, Shanghai Korain Biotech Co. Ltd.); DPP-4 (Cat. No.-E1584Mo, Shanghai Korain Biotech Co. Ltd.) were quantified in serum and BAT as per the manufacturer’s instructions. Analyte concentration in serum is represented as nmol/L (in case of EPI) and ng/L, and in BAT lysate, analyte concentrations were normalized to the tissue weight and expressed as nmol (in case of EPI) and ng/mg of tissue.

## 3. Statistical analysis

Data was presented as mean± SEM. Statistical analysis was performed using GraphPad Prism 8 software (GraphPad, California, United States). Two group comparisons were done using paired or unpaired t-test. Comparison between three or more groups was done with ANOVA (one-way or two-way) followed by Tukey’s post-hoc test. P ≤ 0.05 is considered statistically significant. Specific statistical tests used are mentioned in each figure legend.

## 4. Results

### 4.1 Short-term application of menthol increases CBT, elicits sympathetic and sensory nerves, and BAT activation molecular phenotype

We performed CBT measurement and infrared thermography respectively after 3 days of menthol application **(Fig. 1A).** Menthol induced a significant increase in CBT at 30, 60, and 120 min post-topical application **(Fig. 1B)**. Concurrently, infrared imaging of BAT at the same time points revealed increased heat production or thermogenesis in regions of interest (intrascapular and thoracic) **(Fig. 1C and 1D)**. BAT is densely innervated by sympathetic and sensory nerve fibers, which are activated upon exposure to cold stimuli and release mediators that drive its energy-expending phenotype. To determine whether short-term menthol application elicits this neurochemical activation, we analysed neuronal markers of sympathetic (EPI, ATP, and NPY) and sensory (CGRP and SP) nerves in serum and BAT. Short-term menthol application significantly elevated EPI, ATP, CGRP, and SP levels in both serum **(Supp. Fig. 1A, 1C, 1D, and 1E)** and BAT **(Fig. 1E, IG, 1H, and 1I)**. However, NPY (**Fig. 1F**) was significantly increased only in BAT. These findings indicate that short-term menthol exposure activates both sympathetic and sensory nerve signalling in BAT.

**Figure 1:**
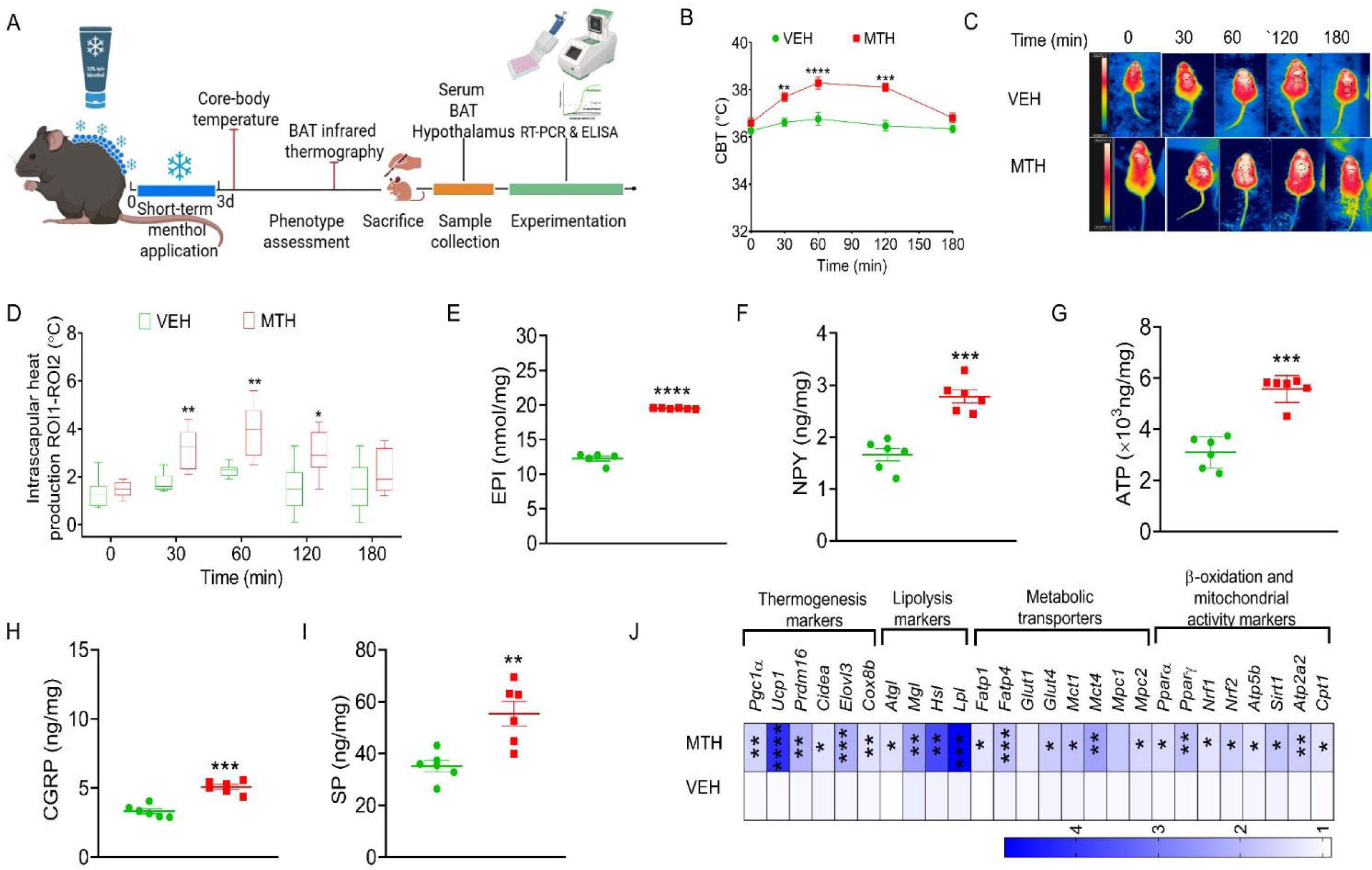
Effect of short-term topical menthol application on CBT, BAT temperature, neuronal markers of sympathetic and sensory nerves and molecular markers of BAT activation. (A) Schematic overview of the experimental design, (B) CBT before (0 min) and after a short-term topical menthol application at 30, 60,120 and 180 min (n=8), (C and D) Representative infrared thermographic images and BAT heat production before (0 min) and after topical menthol application at 30, 60,120 and 180 min (n=6), (E-I) Concentration of EPI, NPY and ATP (SNS markers), and CGRP and SP (Sensory neuronal markers) in BAT (n=6), (J) Heatmap showing relative expression overview of genes in BAT post-menthol (3 day) application (n=6). Data is represented as mean±SEM, analyzed by Student t-test for two-group comparison and Two-way ANOVA for multiple-group comparison, followed by Tukey’s test. p value * < 0.05, ** < 0.01, *** < 0.001, and **** < 0.0001 when compared to VEH group.

We performed gene expression studies for markers of thermogenesis, lipolysis, [-oxidation, and mitochondrial biogenesis. Thermogenesis markers, *i.e., PGC1*_α_*, UCP1*, *PRDM16*, *Elovl3, CIDEA*, *COX8b*, were significantly upregulated post-menthol (3-day) application. Next, lipolysis-related genes like *ATGL,* HSL, *MGL*, and *LPL* were also upregulated in the activated BAT, indicating lipolysis **(Fig. 1J).** Further, mitochondrial biogenesis gene signatures, such as *NRF1*, *NRF2, ATP2B*, *ATP5B,* and *SIRT1,* were also significantly upregulated. We observed significant increase in the gene expression of fatty acid transporters *FATP1* and *FATP4*, and PPARs, *PPAR*_α_ and *PPAR*_γ_, after short-term application of menthol **(Fig. 1J)**. mRNA expression of glucose and lactate transporters (*GLUT4*, *MCT1*, *MCT4*), and mitochondrial pyruvate carrier (*MPC2*) was upregulated, indicating enhanced substrate uptake and utilisation during periods of increased CBT and BAT thermogenesis **(Fig. 1J)**.

### 4.2 Sympathetic and sensory denervation impairs thermoregulation and alters basal physiology

The SNS and sensory system are well known to play a critical role in maintaining energy homeostasis in basal conditions, and recent literature suggests the interdependency [13, 24, 34–36]. For sympathetic denervation, we employed 6-OHDA-based ablation of sympathetic nerves *via* systemic and localized 6-OHDA injections, targeting global and BAT-specific SNS ablation, respectively (**Supp. Fig. 2A)**. Denervation was confirmed by ELISA-based quantification of canonical sympathetic mediator, EPI, in serum and BAT of 6-OHDA-injected mice housed at normal physiological conditions (23°C) and at 4°C for 2 h **(Supp. Fig. 2B – 2E).** Next, for sensory nerve ablation, we employed systemic and BAT-specific denervation of TRPV1-positive sensory neurons using capsaicin (**Supp. Fig. 2F)**. Extent of sensory denervation was quantified by ELISA-based estimation of CGRP in serum and BAT of mice housed at normal physiological conditions and at 4°C for 2 h **(Supp. Fig. 2G-2J).** TRPV1-positive neurons play a role in thermal perception and glucose homeostasis [45, 46]. To assess these effects, an OGTT **(Supp. Fig. 2K and 2L)** and a hot plate test were performed **(Supp. Fig. 2M and 2N)**. The absence of TRPV1 impairs both glucose metabolism and heat sensitivity.

**Figure 2:**
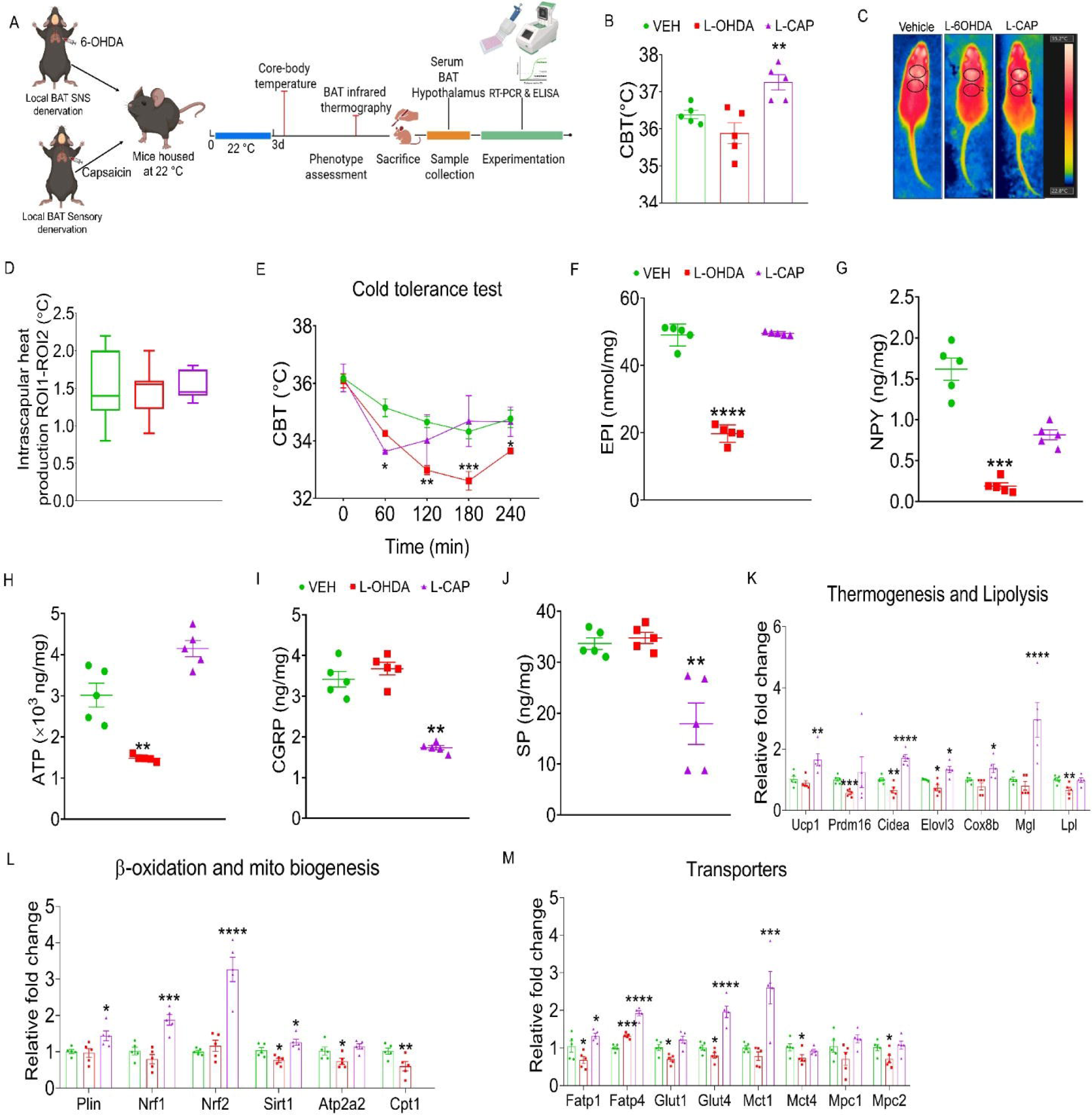
Effect of SNS and sensory denervation on temperature regulation and BAT thermogenesis, and energy homeostasis at normal physiological conditions. (A) Schematic overview of the experimental design, (B) CBT of mice at normal housing conditions (22°C) (n=5); (C and D) Representative BAT infrared thermographic images and BAT heat production at normal housing conditions (24-25°C) (n=5); (E) Cold tolerance test (n=5); (F-J) Serum concentrations of SNS markers-EPI, NPY and ATP (n=5), and sensory neuronal markers-CGRP and SP (n=5) in BAT; (K-M) Relative mRNA expression in BAT for markers of thermogenesis, lipolysis, mitochondrial biogenesis and metabolic transporters under normal physiological conditions (n=5). Data is represented as mean±SEM, analyzed by One-way ANOVA followed by Tukey’s test. p value * < 0.05, ** < 0.01, *** < 0.001, and **** < 0.0001 when compared to VEH group.

To dissect the contribution of sympathetic and sensory innervation to BAT function, we performed BAT-specific denervation. There was no significant change in core temperature and BAT heat production in SNS denervated mice; however, in BAT-specific sensory denervated mice, there was significant increase in CBT, but BAT heat production was not changed **(Fig. 2B-D)**. Next, we conducted cold tolerance test in mice. BAT-specific SNS-denervated mice showed significantly compromised thermoregulation for 4h, whereas BAT-specific sensory nerve ablated mice showed rebound in CBT after 1h and maintained their basal temperature, suggesting the dominant role of SNS in acute cold defence **(Fig. 2E)**. Next, we compared sympathetic and sensory neuronal markers in serum and BAT. In serum, SNS-denervated mice displayed significantly reduced NPY **(Supp. Fig. 1G)** and elevated CGRP and SP levels **(Supp. Fig. 1I and IJ)**, while sensory-denervated mice showed significant reductions in CGRP and SP levels **(Supp. Fig. 1I and IJ)**. In BAT-specific SNS-denervated mice, sympathetic markers (EPI, NPY, and ATP) were significantly decreased, but no effect on sensory markers. Conversely, sensory-denervated mice showed significantly reduced sensory neuronal markers (CGRP and SP), with no significant alterations in sympathetic neuronal markers (EPI, NPY, and ATP) **(Fig. 2F-2J)**. Gene expression analysis revealed that BAT-targeted SNS-denervation led to compromised thermogenesis (*PRDM16, CIDEA*, and *ELOVL3*), whereas sensory denervation resulted in significant upregulation in markers of thermogenesis (*UCP1, CIDEA, ELOVL3*, and *COX8B*) **(Fig. 2K)**. Next, the lipolytic gene, LPL, was downregulated in SNS-denervated mice, and *MGL* was found significantly upregulated in sensory nerve-ablated BAT of mice **(Fig. 2K)**. Mitochondrial biogenesis markers (*NRF1, NRF2, and SIRT1)* were found significantly higher in sensory nerve-ablated mice, whereas *SIRT1, ATP2A2, CPT1*and *PLIN1* were significantly downregulated in SNS-denervated mice **(Fig. 2L)**. Transporters such as *FATP1, GLUT1, GLUT4, MCT4, and MPC2* are significantly downregulated in SNS-enervated mice BAT, whereas *FATP1, FATP4, GLUT4, and MCT1* are significantly higher in sensory nerve-ablated mice **(Fig. 2M)**.

Systemic sympathetic denervation significantly reduced CBT and BAT thermogenesis under standard housing conditions (23 °C), whereas sensory denervation had no significant effect **(Supp. Fig. 3B and 3C, 3D)**. In a cold tolerance test (16 h fasted, 4 h exposure), both sympathetic and sensory nerve–ablated mice exhibited sustained reductions in CBT compared to controls. However, the decline was less pronounced in sensory-ablated mice, indicating partial preservation of thermoregulatory capacity **(Supp. Fig. 3E)**. To assess the impact of sympathetic and sensory denervation on neurochemical signaling, we quantified sympathetic and sensory neuronal markers by ELISA. Sympathetic markers (EPI, NPY, and ATP) were significantly reduced in both serum **(Supp. Fig. 3F-3H)** and BAT **(Supp. Fig. 3K-3M)** of SNS-denervated mice. In sensory-ablated mice, EPI remained unchanged **(Supp. Fig. 3F & 3K)**, NPY was decreased in both serum **(Supp. Fig. 3G)**, and BAT **(Supp. Fig. 3L)**, and ATP was elevated selectively in BAT **(Supp. Fig. 3M)**. Sensory neuropeptide, CGRP was decreased in serum **(Supp. Fig. 3I)** but unchanged in BAT **(Supp. Fig. 3N)** following SNS ablation, whereas significantly reduced in serum **(Supp. Fig. 3I)** and BAT **(Supp. Fig. 3N)** of sensory-ablated mice. Interestingly, SP another neuropeptide significantly increased in serum **(Supp. Fig. 3J)** but remained unchanged in BAT **(Supp. Fig. 3O)** of SNS-ablated mice. In sensory ablated mice, levels of SP significantly reduced in serum **(Supp. Fig. 3J)** and BAT **(Supp. Fig. 3O).**

**Figure 3:**
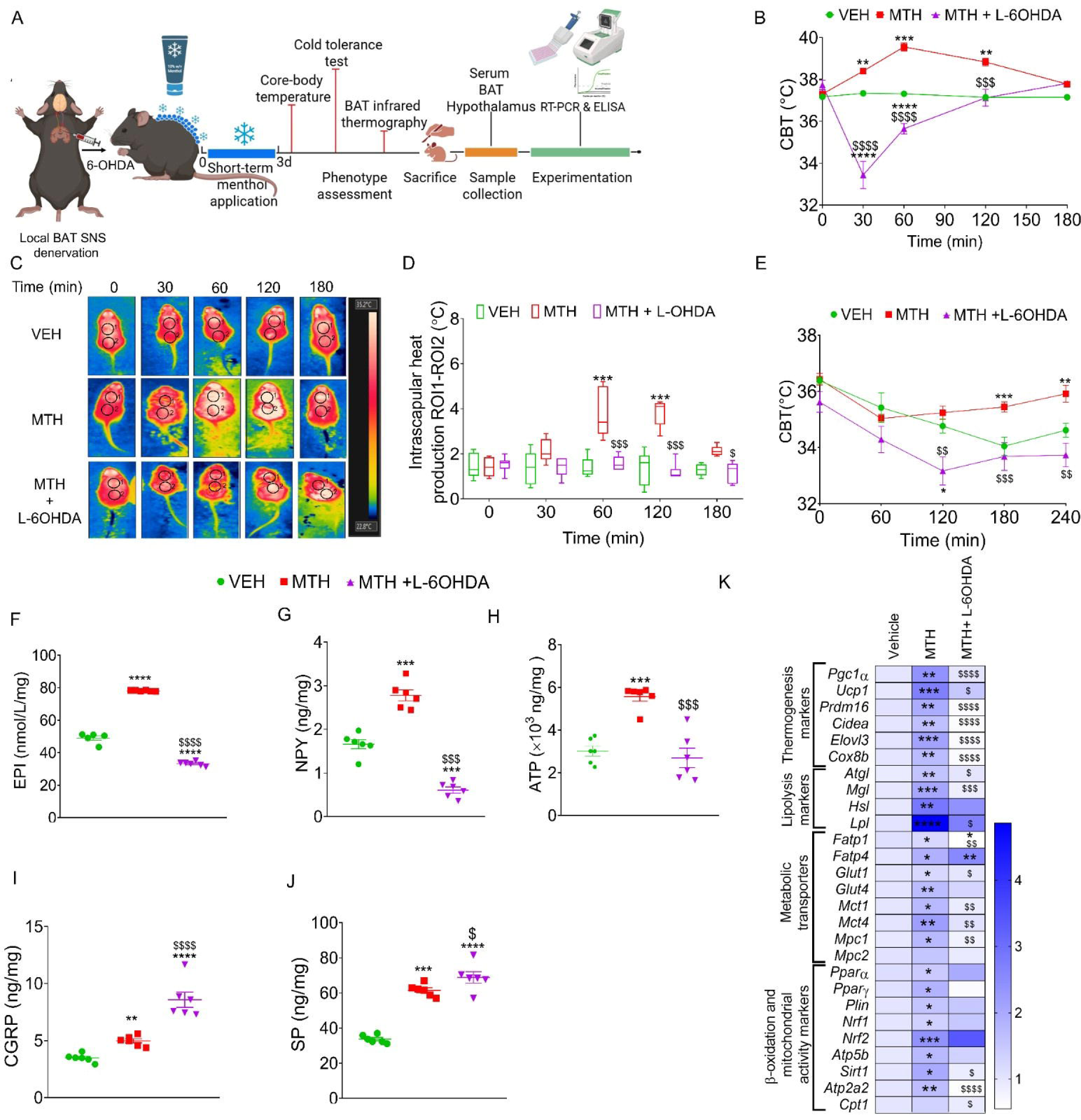
BAT-specific chemical ablation of SNS downregulates CBT, diminishes the BAT thermogenic and energy expanding phenotype after short-term topical menthol application. (A) Schematic illustration of the experimental workflow, (B) CBT before (0 min) and at 30, 60, 120 and 180 min after short-term topical MTH application (n=6-8); (C and D) BAT infrared thermographic images and BAT heat production before (0 min) and after topical menthol application at 30, 60,120 and 180 min (n=6); (E) Cold tolerance test, (n=5); (F-J) Levels of EPI, NPY and ATP (n=6) and CGRP, SP (Sensory neuronal markers) (n=6) in BAT; (K) Heatmap showing relative gene expression BAT (n=6). Data is represented as mean±SEM, analyzed by One-way ANOVA and Two-way ANOVA for multiple groups comparison followed by Tukey’s test. p value * < 0.05, ** < 0.01, *** < 0.001 and **** < 0.0001 when compared to VEH group, ^$^ < 0.05, ^$$^ < 0.01, ^$$$^ < 0.001 and ^$$$$^ < 0.0001 when compared to MTH group.

In SNS-ablated mice, thermogenic genes (*PGC-1*_α_, *CIDEA*, *ELOVL3*, *COX8B*), lipolytic enzymes (*ATGL*, *HSL*, *MGL*), β-oxidation regulators (*PPAR*_α_ and *CPT1*), and mitochondrial biogenesis factors (*NRF2*, *SIRT1,* and *ATP5B*) were broadly downregulated, while *LPL* was upregulated (**Supp. Fig. 3P-3S)**. Sensory ablation induced *CIDEA* and *MGL* upregulation, enhanced mitochondrial biogenesis (*NRF2*, *SIRT1,* and ATP*5B*), and increased expression of *FATP4*, *GLUT1*, and *GLUT4* (**Supp. Fig. 3P-3S)**.

### 4.3 Sympathetic innervations are necessary to regulate CBT and BAT activation and energy-expending phenotype on short-term menthol application

SNS plays central role in thermoregulation and energy expenditure in mammals. Next, to determine whether SNS innervation deficiency in BAT affects CBT, thermogenesis, and whole-body energy expenditure, we performed local chemical denervation of the SNS in BAT by injecting 6-OHDA into both BAT lobes **(Fig. 3A)**. We observed significant drop in CBT post-menthol application in SNS-ablated mice compared to normal mice **(Fig. 3B)**. The local BAT SNS-denervated mice took longer to maintain the basal CBT. BAT infrared thermography also showed attenuation of BAT thermogenesis up to 3 h post-menthol application **(Fig. 3C and D)**. Next, we performed cold tolerance test. The SNS-deneravated mice with menthol application showed significant drop in CBT up to 2 h and shows some rebound in dropped CBT at 3h and 4 h that but unable to maintain their basal CBT during 4h **(Fig. 3E)**. Menthol treated mice CBT decreased up to 1 h, and then there was significant increase in CBT as compared to vehicle-treated mice **(Fig. 3E)**. These results points that sympathetic innervations are necessary for thermoregulation and BAT thermogenesis or heat production (which insulates the body) in response to menthol-induced pharmacological cold. Next, we quantified sympathetic (EPI, NPY, and ATP) and sensory (CGRP and SP) neuronal mediators in serum and BAT. 6-OHDA-induced sympathetic denervation prevented menthol-induced increase in EPI, NPY, and ATP **(Fig. 3F-H)**. In addition, 6-OHDA-induced sympathetic denervation further increased the levels of sensory mediators (CGRP and NPY) in BAT **(Fig. 3I-J)**. This suggests that in the absence of SNS input, sensory pathways may partially compensate to modulate BAT physiology.

Gene expression profiling of menthol treated SNS-denervated BAT revealed that BAT SNS-denervation prevented the menthol induced increase in markers of thermogenesis (*PGC1*_α_*, UCP1, PRDM16, Elovl3, CIDEA* and *COX8B*), lipolysis (*ATGL, MGL* and *LPL*), mitochondrial biogenesis (*SIRT1, ATP2A2,* and *CPT1*), as well as metabolic transporters such as fat transporter (*FATP1*), glucose transporter (*GLUT1*), monocarboxylate transporters (*MCT1* and *MCT4*) and mitochondrial pyruvate carrier (*MPC*1) compared to short-term menthol treated normal mice **(Fig. 3K)**.

Systemic ablation of SNS resulted in a significant reduction in CBT following topical menthol application **(Supp. Fig. 4B)**, and BAT thermography also showed pronounced suppression of BAT thermogenesis in response to topical menthol treatment **(Supp. Fig. 4C and 4D).** Next, in cold tolerance test, showed a significant reduction in CBT as compared to menthol and vehicle-treated normal mice **(Supp. 4E)**. SNS-ablated mice took longer to recover basal core temperature. These findings indicate that loss of sympathetic innervation and BAT thermogenic capacity severely compromises thermoregulatory or core body temperature homeostasis. Consistent with impaired sympathetic signaling, serum and BAT levels of canonical SNS mediators (EPI, NPY, and ATP) were significantly decreased in serum **(Supp. Fig. 4F-4H)** and BAT **(Supp. Fig. 4K-4M)** of complete SNS-ablated mice compared to vehicle-treated normal mice, and menthol treatment in SNS-ablated mice was unable to elevate levels of these mediators. In contrast, sensory neuropeptides (CGRP and SP) were significantly elevated in both serum **(Supp. Fig. 4I and 4J)** and BAT **(Supp. Fig. 4N-4O)**, suggesting compensatory hyperactivation of sensory pathways in response to compromised thermoregulation. Gene expression profiling of BAT from SNS-ablated mice revealed significant downregulation of markers of thermogenesis (*PGC1*_α_*, UCP1, PRDM16, ELOVL3, CIDEA, and COX8b*), lipolysis (*ATGL* and *MGL,* and *LPL*), and mitochondrial biogenesis (*SIRT1, ATP2A2, and CPT1*). Further, transporters such as fat (*FATP1*), glucose (*GLUT1* and *GLUT4*), monocarboxylate (*MCT1* and *MCT4*) transporters, and mitochondrial pyruvate carrier (*MPC1* and *MPC2*) were also significantly downregulated in global SNS-ablated mice **(Supp. Fig. 4P)**.

**Figure 4:**
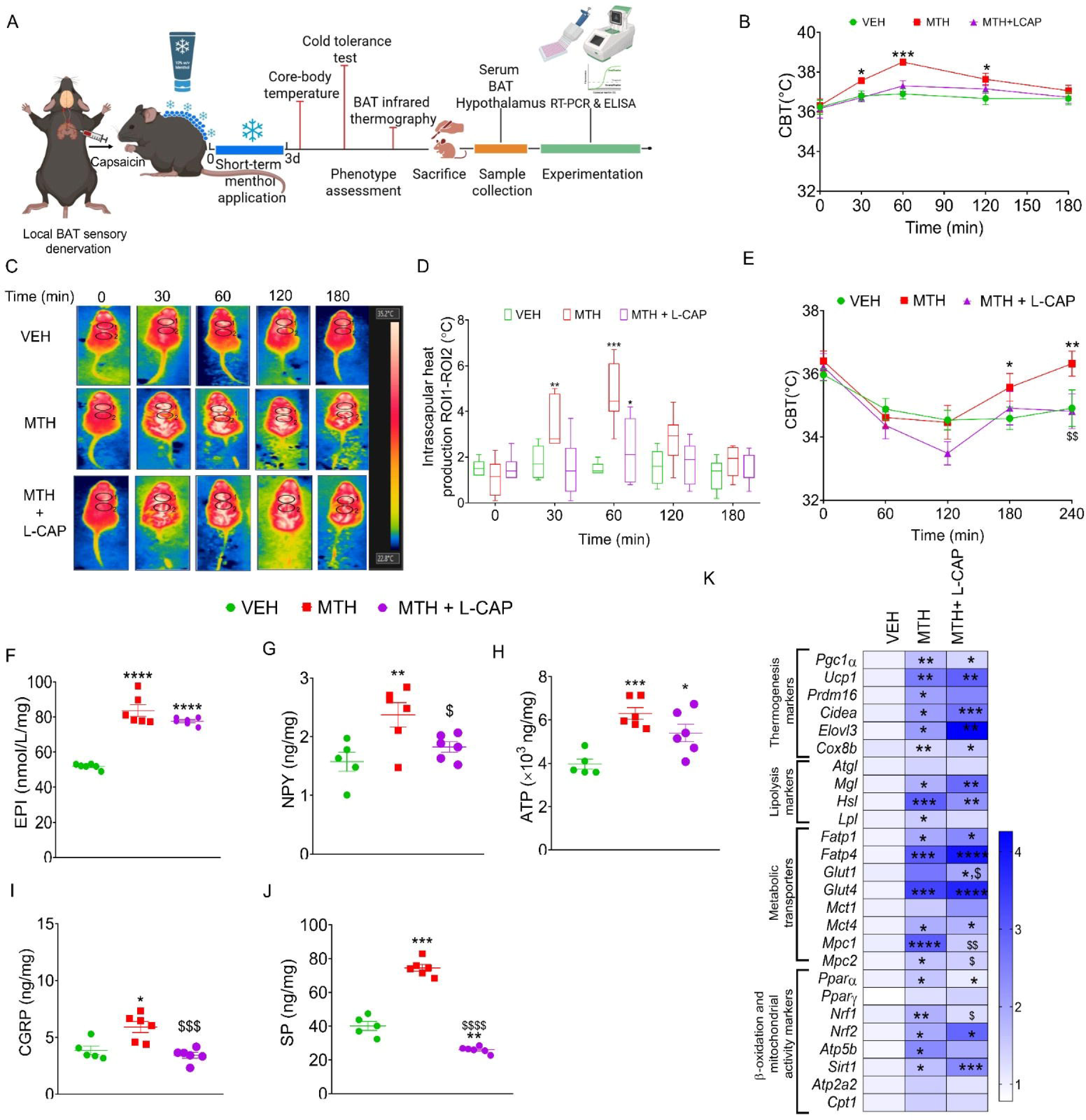
BAT-specific chemical ablation of sensory neurons attenuates topical menthol induced increase in CBT and BAT heat production and partly affects energy expending effects. (A) Schematic overview of the experimental design, (B) CBT measured at baseline (0 min) and at 30, 60, 120 and 180 min post topical menthol (3 day) application (n=6-8) (C and D) Representative infrared thermographic images and quantification of BAT heat production before (0 min) and after topical menthol application at 30, 60,120 and 180 min (n=5-6); (E) Cold tolerance test (n=5); (F-J) BAT concentration of EPI, NPY and ATP and CGRP, SP (n=6); (K) Heatmap of relative gene expression in BAT (n=6). Data is represented as mean±SEM. Statistical significance was assessed using One-way ANOVA and Two-way ANOVA for multiple groups comparison followed by Tukey’s test. p value * < 0.05, ** < 0.01, *** < 0.001 and **** < 0.0001 when compared to VEH group, ^$^ < 0.05, ^$$^ < 0.01, ^$$$^ < 0.001 and ^$$$$^ < 0.0001 when compared to MTH group.

### 4.4 Sensory innervations are not pivotal but partly play role in thermoregulation and BAT activation and thermogenesis on short-term menthol application

BAT is innervated by both efferent sympathetic and afferent sensory neurons, forming a bidirectional feedback loop with CNS to regulate peripheral thermogenic responses. While the role of sympathetic innervation in BAT activation is well established, the contribution of sensory neurons—particularly in response to pharmacological cold mimetics such as menthol—remains poorly understood. To further dissect the role of local sensory innervation, BAT-specific sensory denervation was performed via capsaicin injection into both BAT lobes **(Fig. 4A)**. Menthol-induced increase in CBT and BAT thermogenesis was prevented in BAT-specific sensory nerve-ablated mice **(Fig. 4B-D)**. Next, in cold tolerance test, menthol-induced recovery was prevented in BAT-specific sensory nerve-ablated mice **(Fig. 4E)**. Next, we measured sympathetic nerve mediators (EPI, ATP, and NPY) in menthol-treated BAT-specific sensory nerve-ablated mice. Menthol treatment to BAT-specific sensory nerve-ablated mice significantly increased EPI in both serum **(Supp. Fig. 1F)** and BAT **(Fig. 4F),** similar to menthol-treated (3-day) controls. NPY and ATP did not increase in serum **(Supp. Fig. 1G and 1H)** and BAT **(Fig. 4G and 4H)** post-menthol (3-day) application, pointing to the possibility of sensory nerve-mediated facilitation of the sympathetic nerves in BAT tissue. As expected, sensory neuronal markers (CGRP and SP) estimation showed that BAT-specific sensory nerve ablation prevented menthol-induced increase in CGRP and SP in BAT **(Fig. 4I and 4J**). In case of serum, it only prevented CGRP **(Supp. Fig. 1I and 1J).**

Further, we found a significantly increase in markers of thermogenesis (*UCP1, PRDM16, ELOVL3, CIDEA, and COX8b*), lipolysis (*MGL* and *HSL*), mitochondrial biogenesis (*NRF2, SIRT1, ATP2A2, ATP5B* and *CPT1*), along with these metabolic transporters such as fat (FATP1 and FATP4), glucose (*GLUT1* and *GLUT4*), monocarboxylate (*MCT1* and *MCT2*) and PPARs (*PPAR*_α_ and *PPAR*_γ_) in menthol treated BAT specific nerve denervated mice compared to vehicle and menthol treated mice. This indicates that sensory denervation is possibly increasing sympathetic tone **(Fig. 4K)**.

To investigate the role of whole-body sensory nerves, we employed systemic denervation of TRPV1-positive sensory neurons using capsaicin. Systemic sensory denervation was achieved by systemic injection of capsaicin in mice **(Supp. Fig. 5A)**. CBT infrared thermography showed that systemic sensory nerve–ablation in mice blunted menthol-induced hyperthermia **(Supp. Fig. 5B-D)**. Next, in cold tolerance test, systemic sensory nerve-ablation blunted the ability to regulate CBT as compared to menthol-treated mice **(Supp. Fig. 5E)**. Further, neurochemical analysis showed that EPI levels in serum (**Supp. Fig. 5F**) and BAT **(Supp. Fig. 5K)** were comparable to menthol-treated controls, whereas NPY and ATP were not significantly increased in both serum **(Supp. Fig. 5G and 5H)** and BAT **(Supp. Fig. 5L and 5M)**. Sensory neuropeptides (CGRP and SP) levels were found significantly decreased in both serum **(Supp. Fig. 5I and 5J)** and BAT **(Supp. Fig. 5N and 5O)** post-menthol-treated conditions in sensory nerve-ablated mice.

**Figure 5:**
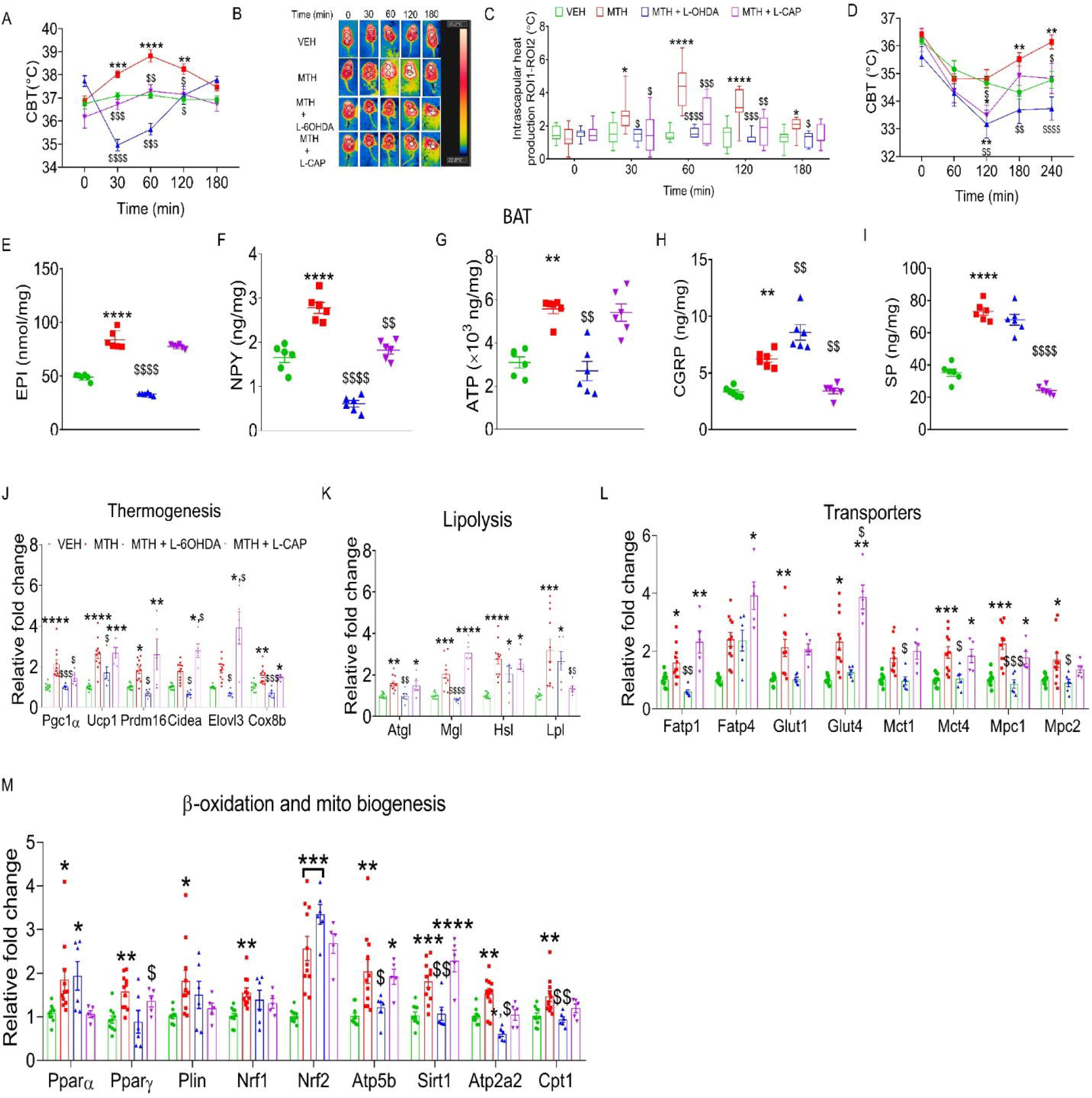
Comparison of short-term menthol-induced effects on thermoregulation and BAT activation and whole body energy expenditure in BAT-specific SNS and sensory denervated. (A) CBT measured at baseline (0 min) and at 30, 60, 120, and 180 min (n=8) ; (B and C) Representative BAT infrared thermographic images and BAT heat production before (0 min) and after topical menthol application at 30, 60, 120 and 180 min (n=6) (D) Cold tolerance test (n=5) (E-I) BAT concentration of SNS markers, EPI, NPY, and ATP (n=6), and sensory neuronal markers-CGRP, and SP (n=6); (J-M) Relative gene expression for markers of thermogenesis, lipolysis, transporters, ℒ-oxidation and mitochondrial biogenesis in BAT (n=6). Data is represented as mean±SEM, analyzed by One-way ANOVA and Two-way ANOVA comparison followed by Tukey’s test. p value * < 0.05, ** < 0.01, *** < 0.001 and **** < 0.0001 when compared to VEH group, ^$^ < 0.05, ^$$^ < 0.01, ^$$$^ < 0.001 and ^$$$$^ < 0.0001 when compared to MTH group.

Gene expression profiling in BAT of sensory-denervated mice revealed that menthol administration to these mice significantly upregulate in marker genes of thermogenesis (*UCP1, PRDM16, CIDEA,* and *COX8B*), lipolysis (*ATGL, MGL,* and *LPL*), mitochondrial biogenesis (*NRF2, SIRT1, ATP5B,* and *CPT1*), along with transporters for fatty acids (*FATP1*and *FATP4*), glucose (*GLUT1* and *GLUT4*), and pyruvate (*MPC1*), and *PPAR*_γ_ **(Supp. Fig. 5P)** as compared to control mice. Interestingly, menthol treated sensory-denervated mice significantly increased *ELOVL3* (thermogenesis marker), *HSL* (Lipolysis marker), *FATP1*(fat transporter), *GLUT-4* (glucose transporter), *MPC1* (mitochondrial pyruvate carrier), and *NRF1* as compared to only menthol-treated mice **(Supp. Fig. 5P)**.

### 4.5 Short-term topical application of pharmacological cold mimetic menthol has contrasting effects on thermoregulation and BAT activity in organ-specific SNS and sensory nerve-ablated mice

Following our individual experiments, we compared the effect of menthol in BAT-specific SNS ablation and BAT-specific sensory ablation. CBT measurement and BAT thermography post-menthol (3-day) application showed that CBT **(Fig. 5A)** and BAT thermogenesis **(Fig. 5B and 5C)** were severely compromised in SNS-denervated mice as compared to sensory denervated mice. CBT in SNS-denervated mice significantly dropped, and even after 2 h, it didn’t reach normal levels. However, in sensory nerve-ablated mice, there is an increase in CBT post-menthol application, but not comparable to menthol-treated normal mice. Similarly, BAT thermogenesis or heat production is significantly decreased up to 3h in BAT-specific SNS-ablated mice, whereas sensory nerve-ablated mice show no significant change in BAT thermogenesis post-menthol (3-day) application. Next, in cold tolerance test, there is drastic decrease in CBT up to 2 h in BAT-specific SNS-ablated mice, mice unable to maintain normal core temperature during test period (4h) **(Fig. 5D)**. In contrast, BAT-specific sensory nerve-ablated mice recover faster and showed an increase in CBT after a drop (up to 2 h). These results clearly show that both SNS and sensory innervations play a role in menthol-induced increase in CBT, and the presence of sympathetic innervation is responsible for keeping the CBT and BAT thermography at basal levels. Next, SNS denervation prevented menthol-induced increase in the serum **(Supp Fig. 1F-1H)** and BAT **(Fig. 5E-5G)** levels of SNS mediators (EPI, NPY, and ATP). However, sensory denervation does not affect menthol-induced increase in EPI and ATP, but it significantly blunted the menthol-induced increase in NPY **(Fig. 5E-5G)**. Interestingly, sensory neuropeptides, CGRP and SP, have different profiles. CGRP levels in serum of SNS-denervated mice are comparable to those of menthol-treated normal mice **(Supp Fig. 1I)**. However, SNS denervation potentiated the menthol-induced increase in SP levels in serum **(Supp Fig. 1J)**. In BAT, there is decrease in menthol-induced increase in CGRP levels, but SP levels are comparable to menthol group. Further, BAT-specific sensory denervation does not affect menthol-induced increase in CGRP and SP levels in serum, whereas it significantly or efficiently prevents menthol-induced increase in these sensory neuronal markers in BAT **(Fig. 5H and 5I)**. Comparison at transcriptional levels shows that menthol-induced thermogenesis (*PGC1*_α_*, UCP1, PRDM16, CIDEA, ELOVL3,* and *COX8B*) **(Fig. 5J),** lipolysis (ATGL and MGL) **(Fig. 5K)**, metabolic substrate transporters (*FATP1*, *GLUT1*, *GLUT4*, *MCT1*, *MCT4*, *MPC1* and *MPC2*) **(Fig. 5L)** and mitochondrial biogenesis (*ATP5B, SIRT1, ATP2A2,* and *CPT1*) **(Fig. 5M)** are significantly downregulated in SNS-denervated mice. Further, BAT-specific sensory denervation has potentiated an increase in thermogenesis (*ELOVL3 and CIDEA*), lipolysis (*MGL*) markers, *PPAR-*_γ_, and transporters such as *FATP1, FATP4, and GLUT4,* suggesting a compensatory enhancement of BAT metabolic activity **(Fig. 5J-5M)**.

### 4.6 Sympathetic and sensory innervations differentially regulate hypothalamic thermosensory ion channels

Hypothalamus in mammals is mainly involved in thermoregulation and controls energy expenditure. Hypothalamus well known to control BAT thermogenesis via various heat-sensing and cold-sensing populations of neurons [47, 48]. We checked the status of heat-sensing and cold-sensing ion channels in the hypothalamus. Short-term menthol treatment in mice showed increased expression of heat-sensing (*TRPV1*, *TRPV2*, *TRPM2*, and *TRPM3*) and cold-sensing (*TRPM8* and *TRPC5*) ion channels. To further dissect the role of BAT sympathetic and sensory innervations, we compared the status of chemosensory ion channels after systemic and BAT-specific sensory and sympathetic nerve ablation. In BAT-specific sensory nerve–ablated mice, *TRPC5*, *TRPV1*, and *TRPM3* were significantly upregulated, whereas *TRPM8* expression was markedly reduced. In contrast, sympathetic nerve ablation under basal physiological conditions resulted in significant downregulation of *TRPM3* and concomitant upregulation of *TRPM8* in the hypothalamus. Importantly, BAT-specific sympathetic denervation abolished the menthol-induced increase in hypothalamic expression of heat-sensing TRP channels (*TRPV1*, *TRPV2*, *TRPM2*, and *TRPM3*). However, cold-sensing channels showed contrasting results: BAT-specific SNS-ablation prevented and aberrated menthol-induced increase in *TRPM8* and *TRPC5* ion channels. These findings demonstrate that BAT sympathetic signalling is essential for menthol-induced BAT thermogenesis, substrate mobilisation, and hypothalamic thermosensory gene activation. Consistent with this, menthol treatment in sensory denervated mice resulted in downregulation of *TRPC5* and *TRPM8*, whereas *TRPV1*, *TRPM2*, and *TRPM3* were significantly upregulated compared to menthol-treated normal mice **(Fig. 6A).**

**Figure 6:**
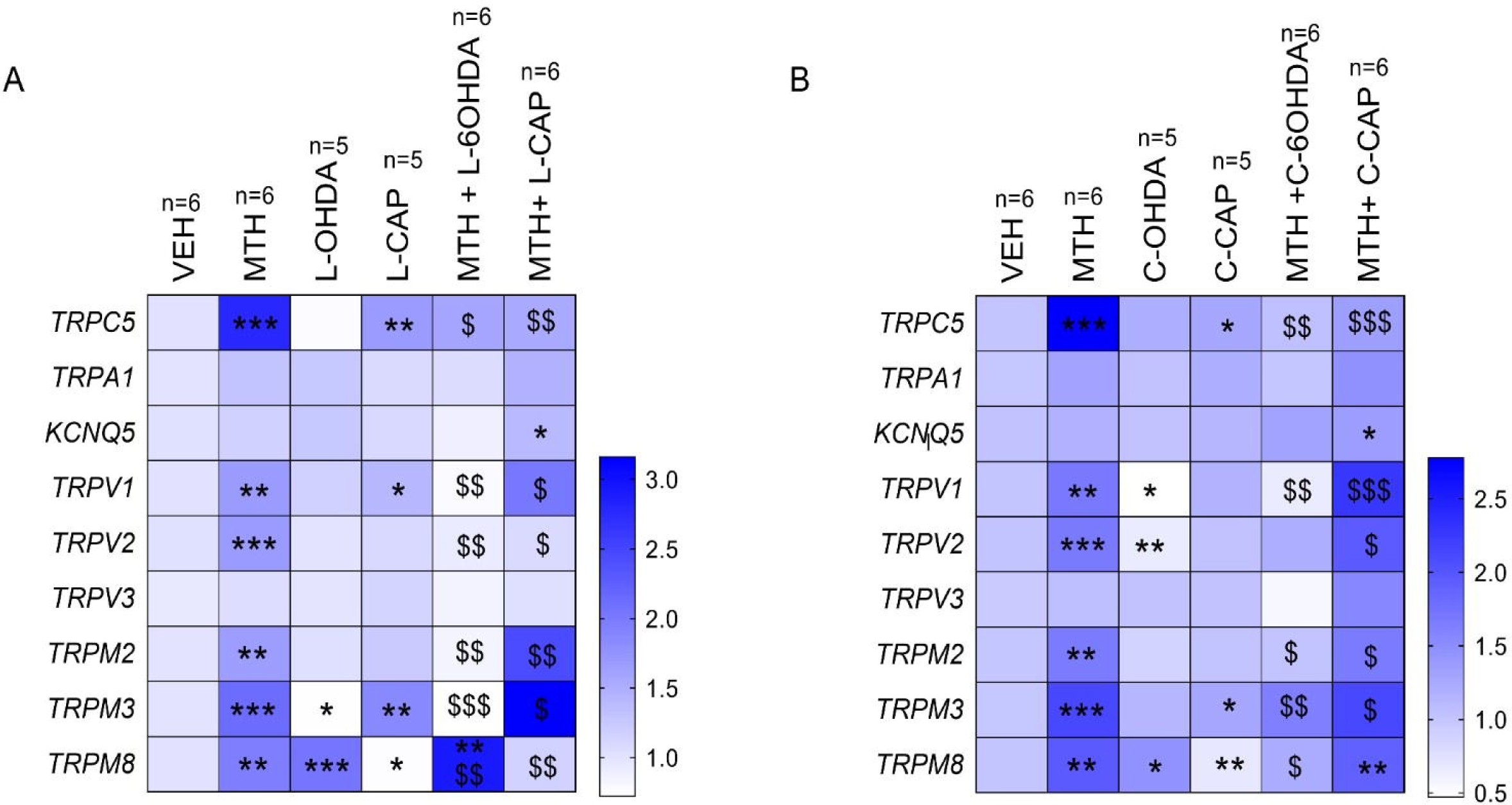
Sympathetic and sensory innervation modulate hypothalamic thermosensory ion channel expression on short-term menthol application. (A) Heat map of relative gene expression of thermos sensing TRP ion channels in hypothalamus post short-term menthol application in BAT-specific SNS and sensory nerve ablated mice (n=5-6); (B) Heat map of relative gene expression of thermosensing TRP ion channels in hypothalamus of systemic SNS and sensory nerve-ablated mice (n=5-6). Data is represented as mean±SEM, analyzed by One-way ANOVA and Two-way ANOVA comparison followed by Tukey’s test. p value * < 0.05, ** < 0.01, and *** < 0.001 when compared to VEH group, ^$^ < 0.05, ^$$^ < 0.01, and ^$$$^ < 0.001 when compared to MTH group.

Further, systemic sympathetic and sensory nerve ablation resulted in distinct expression patterns of hypothalamic thermosensory ion channels under conditions of nerve ablation. In sympathetic nerve–ablated mice, *TRPV1* and *TRPV2* were significantly downregulated, whereas *TRPM8* was significantly upregulated. In contrast, sensory nerve ablation resulted in significant upregulation of *TRPM3* and *TRPC5*, accompanied by downregulation of *TRPM8* in the hypothalamus under normal physiological conditions. Following menthol treatment, systemic SNS ablation led to a significant downregulation of both heat-sensing channels (*TRPV1*, *TRPM2*, and *TRPM3*) and cold-sensing channels (*TRPM8* and *TRPC5*). Conversely, systemic sensory ablation potentiated the menthol-induced upregulation of heat-sensing TRP channels (*TRPV1*, *TRPV2*, *TRPM2*, and *TRPM3*). However, expression levels of cold-sensing channel, *TRPM8* and *TRPC5 i*n menthol treated systemic sensory ablated mice significantly prevented the menthol induced increases observed in normal mice **(Fig. 6B)**. Collectively, our findings underscore the differential contributions of sympathetic and sensory neural circuits in regulating hypothalamic thermosensory ion channel expression and highlight their essential roles in mediating menthol-induced thermogenic signaling.

## 5. Discussion

In this study, we describe the role of sympathetic and sensory neurons that innervate BAT in the acute action of pharmacologic cold mimic, menthol. Notably, using organ-targeted chemical ablation models, we uncovered that activation of BAT thermogenic, lipolytic, and energy-expending program following short-term topical application of menthol (10 %, once a day for 3 days) is dependent primarily but not exclusively on centrally mediated SNS activation. We found that, in addition to SNS neurons, TRPV1-positive sensory neurons may play a significant role, possibly through the release of neuropeptides (CGRP and SP), lipid mediators, and sensitization to temperature/bioavailable menthol, and are also partially necessary for efficient BAT activation. In addition to this, our systemic ablation studies also confirm that both pathways are essential for full BAT activation and energy homeostasis. These findings underscore a dual mechanism where SNS drives and sensory neurons modulate menthol-induced BAT thermogenesis.

Short-term application of menthol, a pharmacologic cold mimic, for 3 simultaneous days significantly increased CBT in a phasic manner. Mice were cold sensitive, and gene expression studies supported energy expending (adaptive thermogenesis, lipolysis, fuel utilization, mitochondrial biogenesis) molecular phenotype in mice’s BAT. In addition, BAT levels of both sensory (CGRP and SP) and sympathetic (NE, NPY, and ATP) neuronal activation markers were significantly higher, pointing to their involvement in the action of menthol. Based on recent findings from multiple groups, it is quite intriguing to hypothesize that there is a feedback loop involving efferent sympathetic input and afferent sensory component to DRGs and CNS, which can work in tandem to maintain homeostasis after menthol-induced changes in BAT [23, 34, 36]. Activation of sensory neurons puts brakes on sympathetic activation and hence maintains thermal and metabolic homeostasis in BAT [24]. To test this hypothesis, we performed systemic and organ-targeted ablation of sympathetic and sensory neurons using chemical methods. One can argue that choosing chemical ablation method made it non-specific ablation, but we strongly believe that genetic ablation of physiologically relevant neurons at young age may have global effects that are compensatory in nature [49, 50], hence using chemical methods in adult mice is justified

As expected, systemic sympathetic neuron ablation prevented menthol-induced increase in CBT. In the absence of sympathetic activation to counter cold, there is prolonged hypothermia in menthol-treated mice. Along with this, the effect of menthol on ELISA markers (in both serum and BAT) and energy-expanding molecular genotype in BAT was prevented in these mice. All this was expected as there is substantial literature supporting the role of the sympathetic system globally and in particular innervations in BAT in the action of cold and cold mimicker, menthol [8, 27, 37, 51–54]. Further, in organ-specific ablation too, there was a decrease in CBT, albeit not up to the extent of systemic ablation, possibly because additional sites, other than BAT, of heat generation, i.e., muscle, WAT, and liver, are still active [55, 56]. In the cold tolerance test, menthol-treated mice were able to develop counterregulatory mechanisms quicker than vehicle mice. We have previously reported that 15 days of topical menthol application makes mice cold-sensitive or warm seekers [20]. However, the extent of compensatory effect of menthol treatment in organ-specific SNS-ablated mice is higher than systemic ablation in mice. Similarly, the sympathetic innervation mediators (NE, NPY, ATP) significantly decreased in ablated mice, both systemic and tissue targeted, which prevented menthol-induced effect. Contrastingly, sensory mediators (CGRP and SP) levels significantly increased in BAT of ablated mice, indicating menthol-induced compensatory increase in sensory signalling. Both systemic and organ-specific sympathetic ablation prevented the effect of menthol-induced thermogenic, fuel-utilizing, and energy-expanding genotype in BAT.

Sensory neurons expressing ion channels such as TRPV1 and Piezo2 serve as key mediators of adipose afferent signaling. A range of physiological inputs—including BAT temperature, lipolytic products, endocrine signals, mechanical stress, and immune mediators (e.g., cytokines and chemokines) either independently or synergistically regulate the sensory afferent activity within BAT [57–59]. Elucidating the roles of these diverse signals is critical for a deeper understanding of the adipose tissue–brain axis, which holds therapeutic potential for metabolic diseases such as obesity. However, recent studies on BAT–brain axis have been conducted under physiological conditions, with limited insight into how this axis operates during pathological states (e.g., obesity, type 2 diabetes) or in response to pharmacological interventions. Addressing this gap is essential before targeting sensory innervation of adipose tissue can be considered a viable strategy for metabolic disease therapy.

We ablated sensory neurons systemically and locally in BAT to understand their role in menthol-induced action in BAT. As expected, systemic ablation raised CBT in mice, but local ablation didn’t increase temperature under normal physiological conditions [60]. Further, the interesting fact is that topical menthol application in both systemic and BAT-ablated mice prevented an increase in CBT. There is a possibility that in case of systemic sensory denervation, there is no scope for further increase in temperature, due to induced hyperthermia. However, this was not the case in organ-targeted ablation, and there too menthol didn’t increase CBT. In case of ELISA, menthol-induced increase in the levels of BAT NPY was prevented in sensory denervated mice. NPY is a potent inhibitor of thermogenesis inhibit local brown adipocyte thermogenesis, decrease in NPY increases BAT thermogenesis as seen in gene expression data.

Menthol administration induces thermogenic and energy-expending phenotype, and sympathetic innervations/signalling play a significant role. Systemic and BAT sensory denervation resulted in significant increase in thermogenic and energy-expending molecular phenotype, which corroborates earlier findings [24, 61] that sensory neurons negatively regulate BAT activation. Gene expression profiling of BAT reveals that sensory neuron ablation amplifies menthol-induced upregulation of several key metabolic genes, including *MGL, GLUT4, FATP4, PPAR*_α_, and notably *ELOVL3.* This suggests that intact sensory innervation modulates the transcriptional or translational regulation of genes central to lipid metabolism, insulin sensitivity, and fatty acid elongation. The enhanced expression of *FABP5* and particularly *ELOVL3* following menthol treatment, with further upregulation after sensory denervation, points toward increased lipogenesis and the potential for lipid futile cycling. This may also relate to observed reductions in *NPY*, as previous studies link NPY signaling to lipid accumulation and BAT whitening [62]. Moreover, increased expression of PPARs, especially *PPAR*_α_ post-denervation, highlights a possible role for sensory nerves in maintaining lipid homeostasis. Overall, our gene expression data is pointing to the significant role played by both systemic and BAT-specific sensory signalling.

Further, menthol administration significantly increases both heat-sensitive (*TRPV1*, *TRPV2*, *TRPM3*, and *TRPM3*) and cold-sensitive (*TRPC5* and *TRPM8*) TRP channels [63–65]. BAT-specific sympathetic and sensory denervation significantly increased the expression of TRPM8 and TRPC5 (also *TRPM3*) respectively, suggesting the regulation of specific cold-sensitive neurons through each innervation. BAT-specific sympathetic denervation prevented menthol-induced changes in hypothalamic gene expression of TRP channels, except TRPM8, which was significantly increased. BAT-specific sensory denervation further increased the expression of heat-sensitive TRP channels (*TRPV1, TRPM2, TRPM3*) but prevented the increase in cold-sensitive TRP channel expression (*TRPC5 and TRPM8*). These results suggest the complexity of heat and cold-sensitive neurons in hypothalamus and their interaction with sympathetic and sensory innervations in BAT, both in physiological and pharmacological (menthol application) conditions. Studies using knockout mice and optogenetic screens will be required to delineate the role of heat and cold-sensitive neurons. Overall, there are many open questions still to be answered, but our work in this paper at least scratched the surface.

This study has two primary limitations. While we employed established protocols for sensory neuron-specific denervation, the potential contribution of non-neuronal TRPV1-expressing cells in BAT remains unresolved. These cell types may independently influence the observed thermogenic responses. Additionally, we did not identify the specific mediators or molecular pathways downstream of sensory denervation that contribute to BAT activation. In summary, this work identifies that both sympathetic and sensory innervations regulate menthol-induced increase in energy-expending molecular phenotype in BAT. This highlights the broader influence of sensory modalities—including temperature, immune/inflammatory cues, and mechanosensation on adipose tissue metabolism. These findings offer a new direction for the development of strategies to prevent or treat obesity through cold-mimicking approaches.

## CRediT authorship contribution statement

Roshan Lal: Data curation, Formal analysis, Investigation, Methodology, Writing – original draft; Neha Soni: Investigation, Methodology, Visualization; Kushhagra Agarwal: Formal analysis, Methodology; Kanthi Kiran Kondepudi: Writing – original draft, Writing – review & editing; Pragyanshu Khare: Writing – review & editing; Katharina Zimmermann: Writing – review & editing, Resources; Małgorzata Karbownik-Lewińska: Writing – original draft, Writing – review & editing; Adam Gesing: Formal analysis, Writing – original draft, Writing – review & editing; Vaishali Aggarwal: Methodology, Formulation of menthol cream; Kanwaljit Chopra: Writing – original draft, Writing – review & editing; Mahendra Bishnoi: Conceptualization, Investigation, Methodology, Project administration, Resources, Supervision, Visualization, Writing – original draft, Writing – review & editing.

## Acknowledgments

The authors acknowledge the funding support provided by BRIC-NABI, Department of Biotechnology, GoI, Department of Science and Technology (DST), GoI, Indian council of medical research (ICMR), GoI (Grant number IIRPIG-2023-0000640), and The Humboldt Foundation Exchange Program to Dr. Mahendra Bishnoi and Prof. Katharina Zimmermann. The authors also acknowledge the support provided by ICMR, GoI, for providing fellowship to Mr. Roshan Lal. Authors also acknowledge UGC, GoI, for providing fellowship to Ms. Neha Soni. We acknowledge the use of artificial intelligence tool, ChatGPT4 to refine the language and for better flow of information in this manuscript.

## Data and materials availability

All data needed to evaluate the conclusions in the paper are present in the paper and/or the Supplementary Materials. Additional data related to this paper may be requested from the authors.

## Supp. Figure legends

**Supp. Figure-1:**
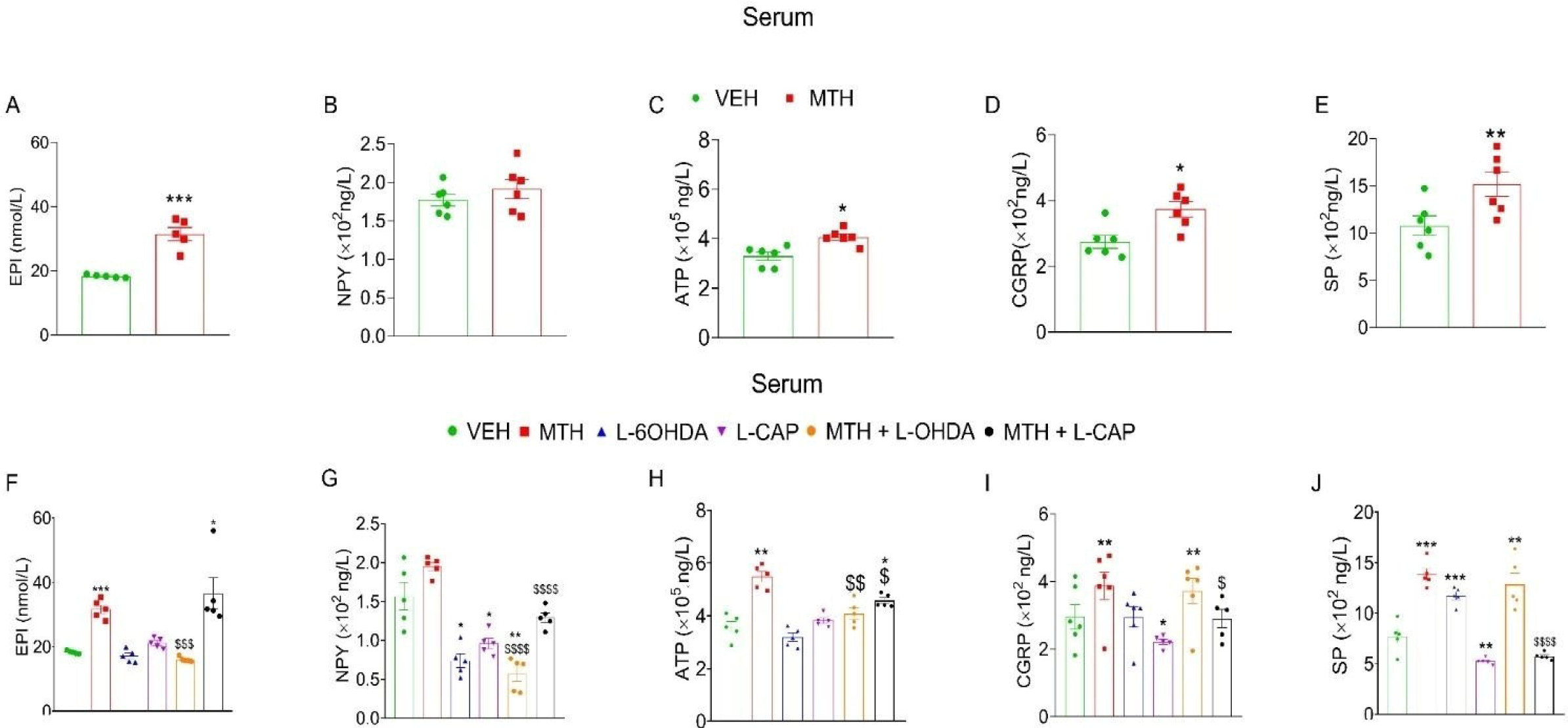
Effect of short-menthol application on SNS and sensory neuronal markers in normal mice and BAT specific SNS and sensory nerve ablated mice. (A-E) Concentration of EPI (A), NPY (B), ATP(C) CGRP (D) and SP (E) in serum (n=5); (F-J) Serum concentration of EPI, NPY, ATP, CGRP and SP (Sensory neuronal markers) post menthol application in BAT-specific SNS and sensory nerve ablated mice (n=5). Data is represented as mean±SEM, analyzed by One-way and Two-way ANOVA followed by Tukey’s test. p value * < 0.05, ** < 0.01, *** < 0.001 and **** < 0.0001 when compared to VEH group, ^$^ < 0.05, ^$$^ < 0.01, ^$$$^ < 0.001 ^$$$$^ < 0.0001 when compared to MTH group.

**Supp. Figure-2:**
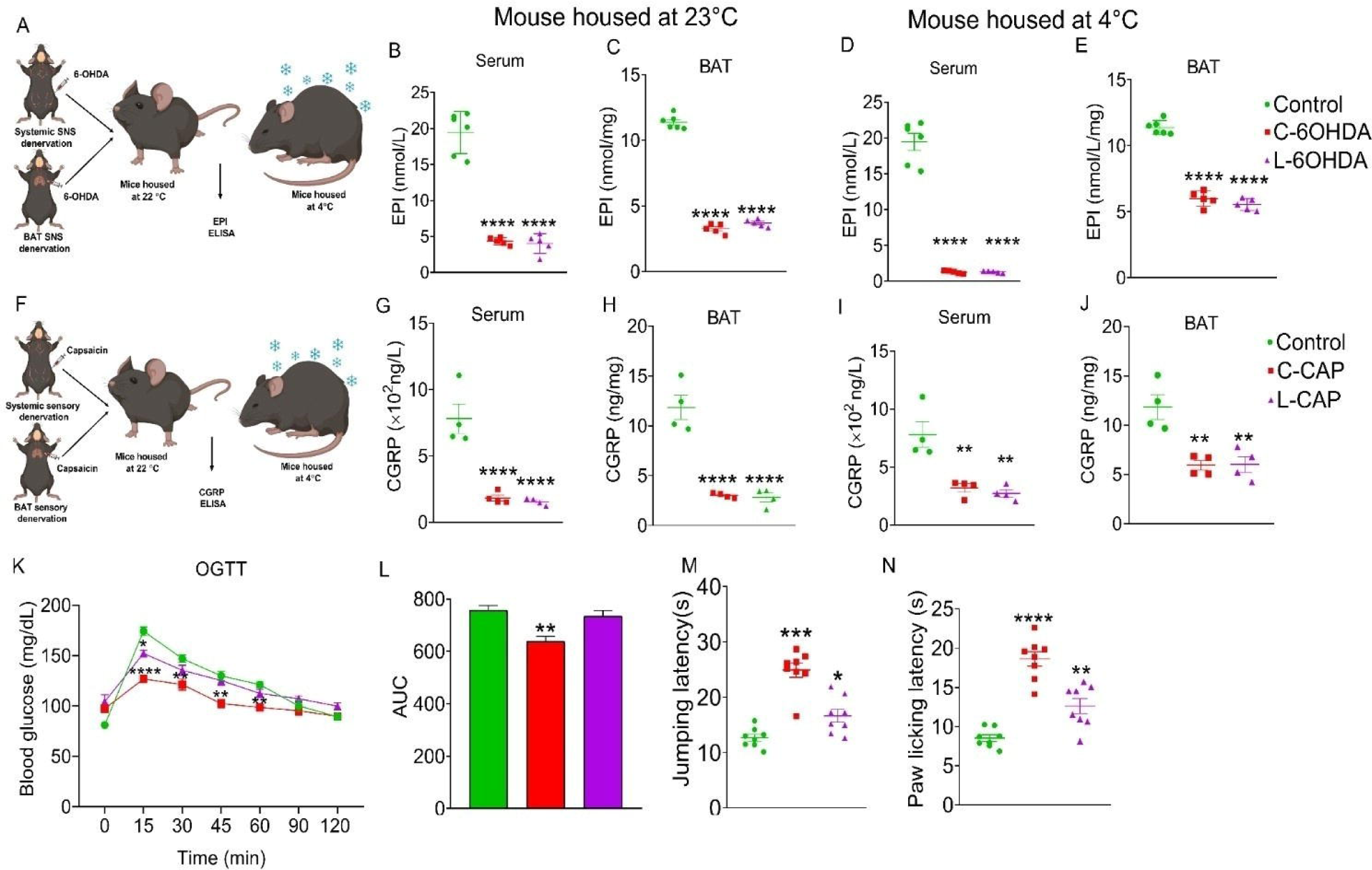
Characterization of Systemic and BAT specific-ablation of SNS and sensory nerves in mice model. (A) Schematic representation of 6-OHDA injection and experiment plan; (B and C) EPI concentration in serum and BAT of mice housed at 22°C (n=5-6); (D and E) ) EPI concentration in serum and BAT of mice housed at 4°C (n=5-6); (F) Schematic of capsaicin injection and experiment plan; (G and H) CGRP concentration in serum and BAT of mice housed at 23°C (n=4-5); (I and J) EPI concentration in serum and BAT of mice housed at 4°C (n=5-6); (K and L) OGTT and AUC (n=8); (M and N) Jumping and paw licking latency in hot plate test (n=8). Data is represented as mean±SEM, analyzed by Student t-test for two-group comparison and Two-way ANOVA for multiple groups comparison followed by Tukey’s test. p value ** < 0.01, *** < 0.001, and **** < 0.0001 when compared to VEH group.

**Supp. Figure 3:**
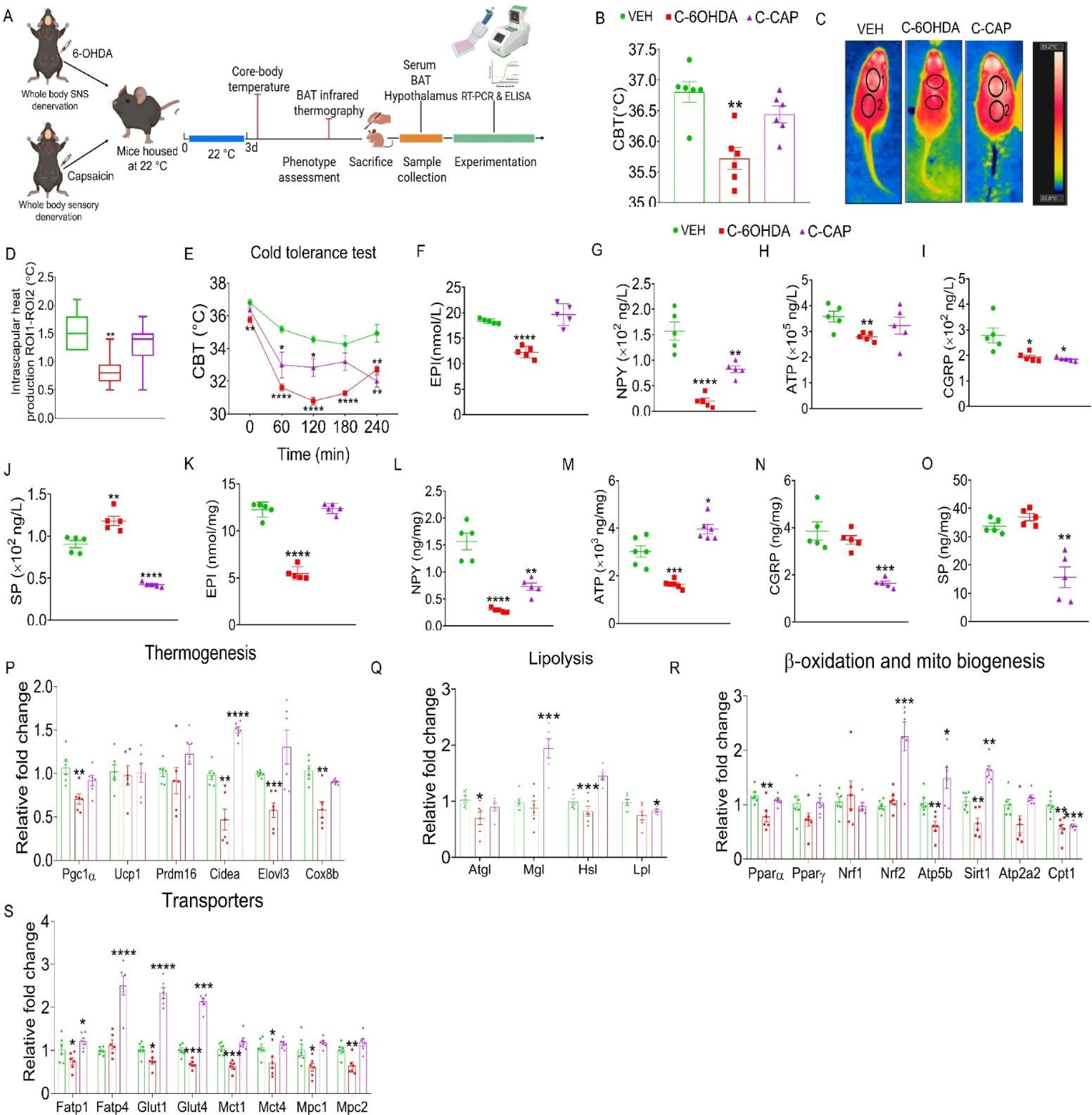
Effect of systemic SNS and sensory denervation on temperature regulation and BAT thermogenesis, and energy homeostasis at normal physiological conditions. (A) Schematic representation of experimental plan; (B) CBT of mice at normal housing conditions (22°C) (n=6); (C and D) Representative BAT infrared thermographic images and BAT heat production at normal housing conditions (22°C) (n=6); (E) Cold tolerance test (n=5); (F-J) Serum concentrations of SNS markers, EPI, NPY and ATP and sensory neuronal markers, CGRP and SP (n=5); (K-O) BAT concentrations of EPI, NPY and ATP, CGRP and SP (n=6); (P-S) Relative mRNA expression in BAT for markers of thermogenesis, lipolysis, mitochondrial biogenesis and metabolic transporters under normal physiological conditions (n=6). Data is represented as mean±SEM, analyzed by One-way ANOVA followed by Tukey’s test. p value * < 0.05, ** < 0.01, *** < 0.001, and **** < 0.0001 when compared to VEH group.

**Supp. Figure-4:**
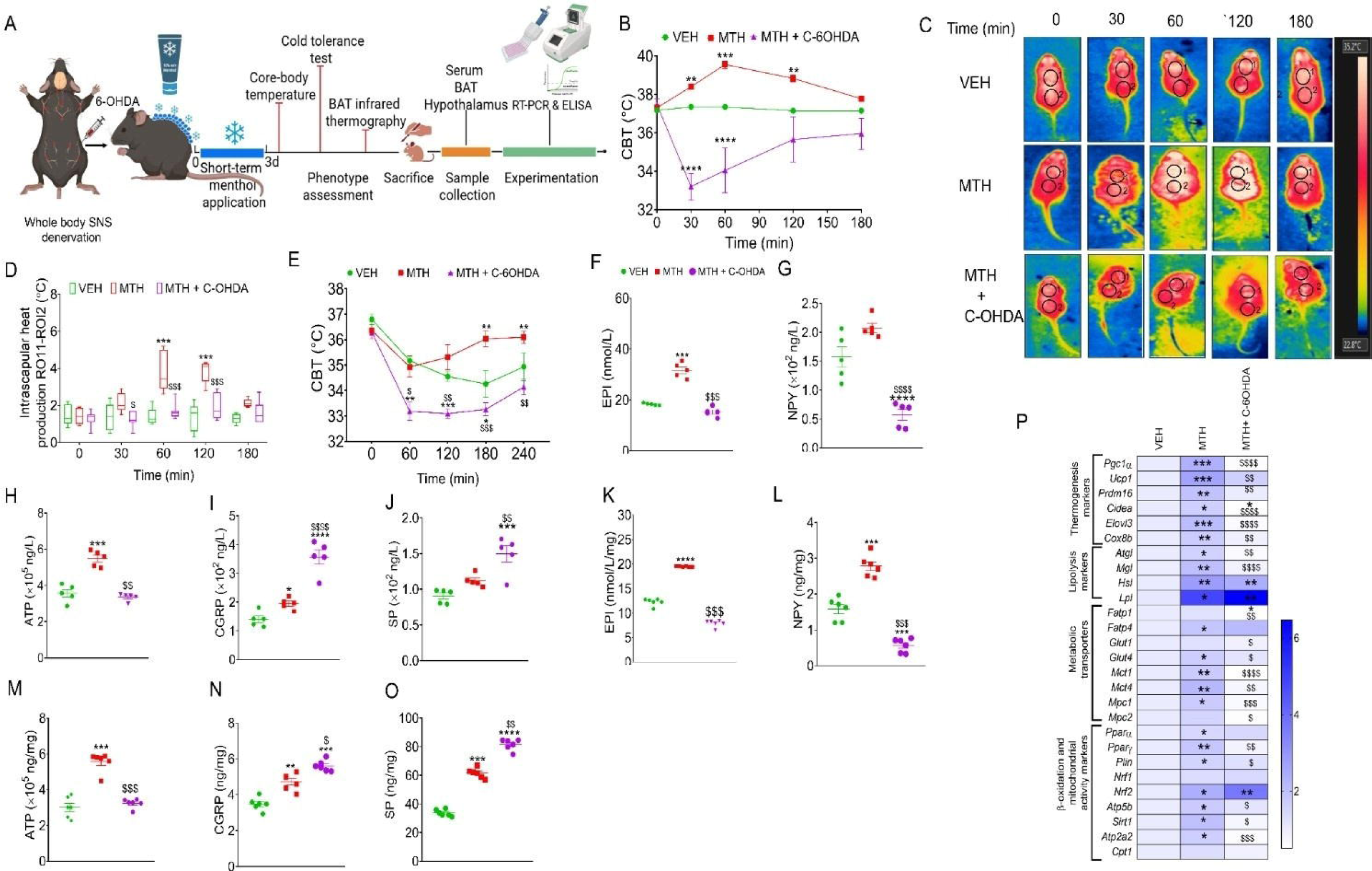
Systemic ablation of whole body SNS compromise thermoregulation, BAT thermogenesis and activation and downregulates energy expending transcription programs. (A) Schematic overview of the experimental workflow, (B) CBT before (0 min) and at 30, 60, 120 and 180 min after short-term topical MTH application (n=6-8); (C and D) Representative BAT infrared thermographic images and heat production before (0 min) and after topical menthol application at 30, 60,120 and 180 min (n=6); (E) Cold tolerance test (n=5); (F-J) EPI, NPY, ATP, CGRP and SP levels in serum (n=5); (K-O) EPI, NPY and ATP, CGRP and SP levels in in BAT (n=6) (P) Heatmap depicting relative expression of thermogenic and metabolic genes in BAT (n=6). Data is represented as mean±SEM, analyzed by One-way and Two-way ANOVA followed by Tukey’s test. p value * < 0.05, ** < 0.01, *** < 0.001 and **** < 0.0001 when compared to VEH group, ^$^ < 0.05, ^$$^ < 0.01, ^$$$^ < 0.001 and ^$$$^ < 0.0001 when compared to MTH group.

**Supp. Figure 5:**
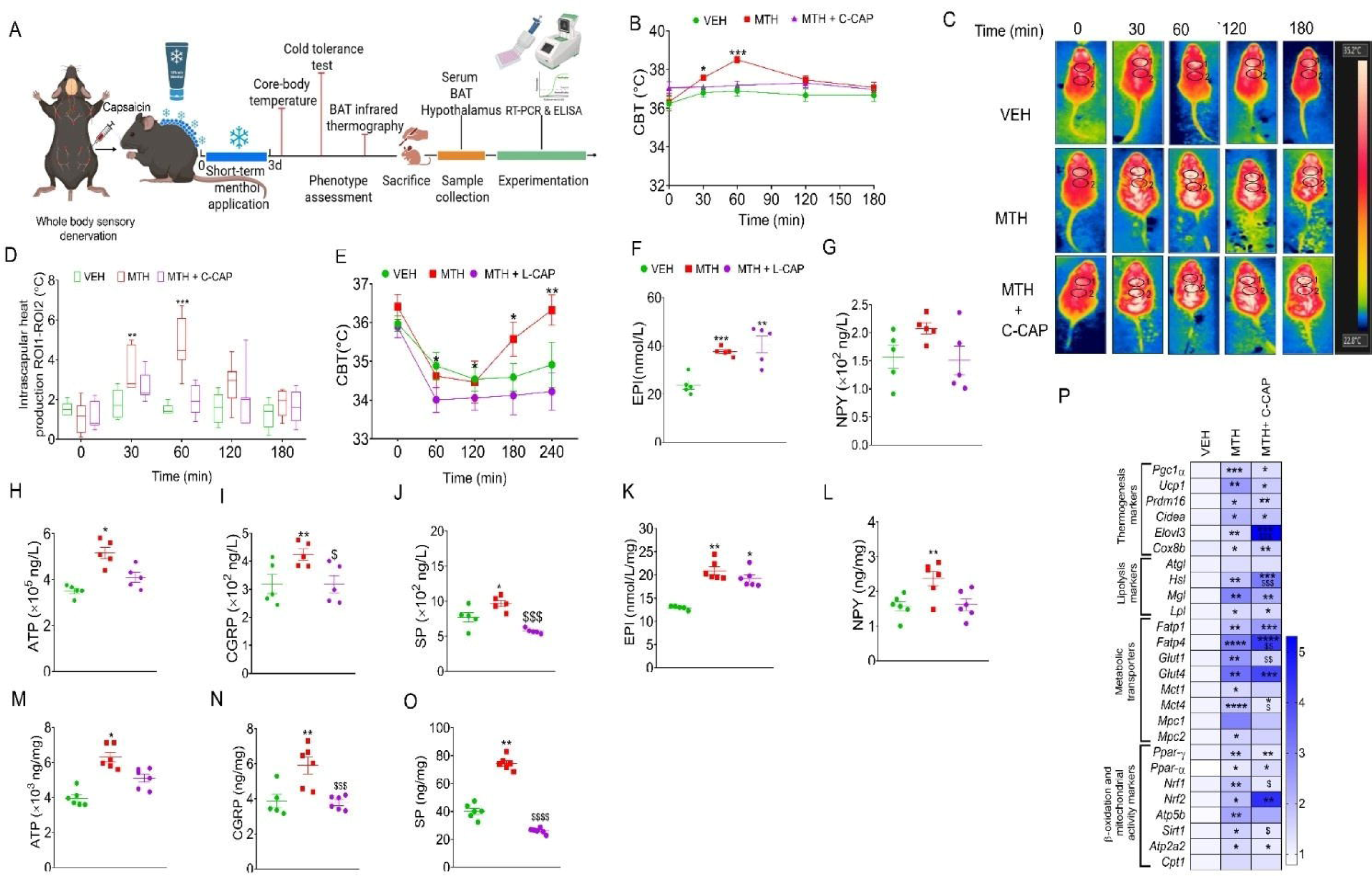
Systemic ablation of whole-body sensory neurons induces hyperthermia and augments menthol-induced BAT thermogenesis and energy-expending effects. (A) Schematic representation of experimental design, (B) CBT measured at baseline (0 min) and at 30, 60, 120 and 180 min following short-term topical MTH application (n=5-8); (C and D) Representative infrared thermographic images of BAT and corresponding BAT heat production before (0 min) and after topical menthol application at 30, 60,120 and 180 min (n=5-8); (E) Cold tolerance test (n=6) (F-J) Serum concentration of EPI, NPY, ATP, CGRP and SP (n=5); (K-O) BAT concentration of EPI, NPY, ATP, CGRP, and SP (n=6); (P) Heatmap showing relative gene expression in BAT (n=6). Data have been represented as mean±SEM, analyzed by One-way and Two-way ANOVA followed by Tukey’s test. p value * < 0.05, ** < 0.01, *** < 0.001 and **** < 0.0001 when compared to VEH group, ^$^ < 0.05, ^$$^ < 0.01, ^$$$^ < 0.001, and ^$$$$^ < 0.0001 when compared to MTH group.

